# Detection and Molecular Epidemiology of Human Bocavirus in Children with Acute Gastroenteritis from Brazil

**DOI:** 10.1101/404632

**Authors:** L.S. Soares, A. B. Lima, K.C. Pantoja, P.S. Lobo, J.F. Cruz, S.F.S. Guerra, D.A.M. Bezerra, R.S. Bandeira, J.D.P. Mascarenhas

## Abstract

Human Bocavirus (HBoV) is a recently discovered virus and was first detected in the nasopharyngeal aspirate samples and after in stool samples, suggesting that HBoV may be a causative agent for human enteric infections. Due to absence of treatment options, there is a need to understand the disease-causing mechanism of these viruses. The aim of this was to demonstrate the prevalence of HBoV from children less than 10 years with acute gastroenteritis in Brazil, during November 2011 to November 2012. Stool samples from hospitalized children ≤ 10 years who presented symptoms of acute gastroenteritis were analyzed for the presence of HBoV DNA by nested-PCR. HBoV- positivity was detected in 24.0% (54/225) of samples. Two peaks of HBoV detection were observed, during November 2011 and July to September 2012. Co-infections between HBoV and rotavirus A were identified in 50.0% (27/54) of specimens. Phylogenetic analysis identified the presence of HBoV-1 (94.8%), HBoV-2 (2.6%) and HBoV-3 (2.6%) species, with only minor variations among them. Further investigations are necessary to improve the knowledge on the role of HBoV in gastrointestinal infections.

## INTRODUCTION

Globally, diarrhoeal disease in 2015 were responsible for 499 000 deaths of children aged < 5 years (1). Several pathogens are associated with acute gastroenteritis (AGE), however ∼40% of cases remain of unknown etiology (2). Human Bocavirus (HBoV) is a recently discovered virus and was first detected in the nasopharyngeal aspirate samples and after in stool samples, suggesting that HBoV may be a causative agent for human enteric infections (3–5).

HBoV belongs to *Parvoviridae* family, *Parvovirinae* subfamily, *Bocaparvovirus* genus (6). This parvovirus is a small single stranded DNA virus with a diameter of 18–26 nanometers and contains a non-enveloped icosahedral capsid (7). The genome of HBoV consists of three ORFs that encode two non structural proteins (NS1 and NP1) and two structural proteins (VP1 and VP2) (7,8). Based on genetic variability of VP1 region, HBoV are divided into 4 species: HBoV1 - HBoV4. Several groups have detected HBoV1 in respiratory tract infections, while HBoV2, 3 and 4 were reported in fecal samples (9,10).

The role of HBoV in respiratory disease and acute gastroenteritis remains unclear. It is believed that HBoV can persist for a longer period in respiratory tract mucosa, reaches the bloodstream or ingestion and migrates to gastrointestinal tract, where it may generate a new infection or be excreted in an asymptomatic way (11). However, there are increasing reports on HBoV detection in fecal specimens from patients with acute diarrhoea, particularly in children (12–14).

HBoV have been detected mostly in respiratory tract secretions and stool, however the viruses have been found in serum and cerebrospinal fluid causing viremia (15,16). Likewise, several studies have reported HBoV in sewage and river water (17– 19).

Since the first HBoV detection, several studies reported its association with respiratory and gastrointestinal infections, mostly in young children (12,20,21). Nevertheless, the significance of this virus as a causative agent in human infections remains unknown, due to high prevalence of co-infection with other viruses in symptomatic patients as well as to frequent detection of HBoV in asymptomatic individuals. A recent systematic review suggests that even though HBoVs are often considered bystanders in acute gastroenteritis course, HBoV-2 infection may increase the risk for this disease (5,10,22).

Several studies have reported HBoV associated with respiratory and gastrointestinal tract infections as single pathogens, causing severe disease (21–23). Due to absence of treatment options, there is a need to understand the disease-causing mechanism of these viruses.

(10) have reported that HBoV prevalence was similar between subjects with or not gastrointestinal symptoms. HBoV has also been found to co-infect humans with other enteric viruses. A study conducted in Pakistan has related a frequency of 98% involved co-infection between HBoV and rotavirus from children with AGE (24). In Brazil in an investigation that also enrolled diarrheic children have described HBoV co-infection with norovirus, adenovirus and rotavirus in 19%, 3% and 3% of patients, respectively (12).

The aim of this was to demonstrate the prevalence of HBoV from children less than 10 years with AGE in Brazil, from November 2011 to November 2012, to report co-infection with rotavirus A (RVA) and to describe HBoV species.

## METHODS

### Ethics Statement

This study was approved by Evandro Chagas Institute’s Human Research Ethics Committee, protocol number 414.389, in accordance with National Health Council’s Resolution 466/2012. The authors ensure that all procedures contributing to this work comply with the ethical standards of the relevant national and institutional committees on human experimentation and with the Helsinki Declaration of 1975, as revised in 2008.

### Clinical Specimens

Samples from this study were collected from hospitalized children ≤ 10 years who presented symptoms of AGE and were selected from four states in Northern region of Brazil. During November 2011 to November 2012 a total of 541 fecal samples was collected and an aliquot of each sample was stored at –20°C and transported to Evandro Chagas Institute, a Brazilian Ministry of Health’s National Rotavirus Reference Laboratory. Acute diarrhoea was defined as the presence of three or more liquid and semi-liquid stools in a 24-hr period for up to 14 days. For the present study, it was selected 225 samples for HBoV testing, considering a 95% confidence level and a sampling error of 5%.

### Detection of RVA

Fecal samples were screened for the presence of RVA by a commercially available enzyme-linked immunosorbent (ELISA) assay according to the manufacturer’s instructions (*Ridascreen* kit, R-Biopharm^®^, Darmstadt, Germany).

### HBoV Screening and DNA Extraction

Viral genome was extracted from 10% fecal suspensions using guanidinium isothiocyanate-silica method (25). Posteriorly, samples were subjected to nested-PCR targeting the partial region of VP1 gene. The first-round of the nested-PCR was carried out using the AK-VP-F1 and AK-VP-R1 primers set, and the second round using the AK-VP-F2 and AK-VP-R2, as described previously (26).

### Nucleotide Sequencing and Phylogenetic Analysis

Sequencing of the PCR amplicons HBoV strains were performed using the same primers as those used in the second-round of the nested-PCR and carried out with a Big Dye Terminator cycle sequencing kit v 3.1 (Applied Biosystems, Foster City, CA). Electrophoresis was performed on the ABI Prism 3130xl automatic sequencer (Applied Biosystems) and the sequences obtained were aligned and edited using the BioEdit Sequence Alignment Editor program (v. 7.0.5.2). Neighbor-joining method was used to perform the phylogenetic analysis, in which distance was calculated from aligned sequences (27). Dendrograms were constructed using MEGA program v.5.0.1, and bootstrap analysis was performed using 2,000 replicas. Partial nucleotide sequences from this study were deposited in the GenBank database (http://www.ncbi.nlm.nih.gov), under the following access number: MH003642-MH003679.

### Statistical Analysis

HBoV frequencies and genotypes description were calculated using Microsoft Excel software. Comparisons of HBoV infection rates in distinct groups were performed using chi-square test (χ^2^) through BioEstat 5.0 with statistical significance established of P values <0.05 (28).

## RESULTS

From November 2011 to November 2012 an overall HBoV-positivity was detected in 24.0% (54/225, range 0-50%) of hospitalized children ≤ 10 years using Nested-PCR. Figure 1 show the monthly frequencies of HBoV detection, with two peaks, where HBoV rates were over 30%, November 2011 and July to September 2012. Samples were tested for RVA which is the most important viral agent associated with acute gastroenteritis. RVA detection was 40.9% (92/225) and it was observed co-infection between HBoV and RVA in 50.0% (27/54) of samples with median age of 17 months (data not shown).

**Figure 1.**
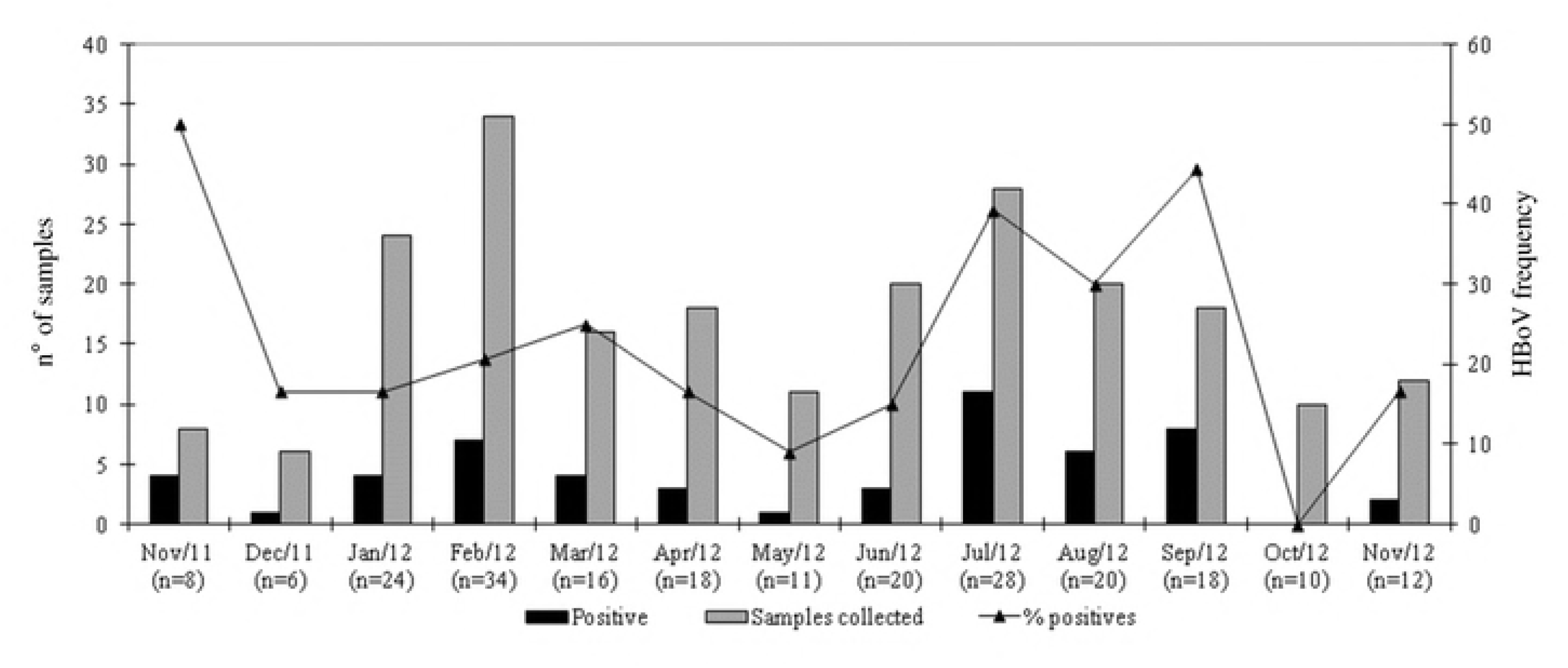
Temporal distribution of HBoV in Brazil, 2011-2012.

General characteristics of HBoV-positive cases are summarized in Table I. The median age of children with HBoV was 16 months, with higher HBoV positivity (66.6%, 36/54) was observed among children aged 7–24 months. With respect to gender, 59.2% (32/54) of infected children were male. No statistically significant differences (P>0.5) were seen when comparing age groups and genders.

**Table I.** Baseline Characteristics of HBoV-Positive Cases in Brazil, 2011-2012.

|  | HBoV positive/<br>tested (%) |
| --- | --- |
| Age group (months) |  |
| 0-6 | 11/50 (22.0) |
| 7-24 | 36/126 (28.6) |
| 25-60 | 6/45 (14.2) |
| >61 | 1/4 (25.0) |
| Clinical characteristics |  |
| Fever | 24/110 (21.8) |
| Vomiting | 28/141 (17.0) |
| Gender |  |
| Male | 32/134 (23.8) |
| Female | 22/91 (24.1) |
| Brazilian State |  |
| Acre | 13/47 (27.6) |
| Amazonas | 38/133 (28.5) |
| Pará | 2/40 (5.0) |
| Roraima | 1/5 (20.0) |

Phylogenetic analysis of partial region of VP1 genes were performed in 38 (70.3%) HBoV positive samples, showing that HBoV1 were identified in 94.8% of samples (n=36), while one isolate each belonged to HBoV-2 and HBoV-3 genotypes, respectively. All HBoV1 strains shared 97–100% and 93.1-100% nucleotide (nt) and amino acid (aa) similarities among themselves, respectively. When compared with another isolates nt and aa similarities ranged from 95.8–100% and 93.2–100%, respectively. With regards HBoV2 and HBoV3 strains showed nt similarities ranged from 93.3–98.6% and 97.1–98.2% with prototype strains, respectively (Figure 2).

**Figure 2.**
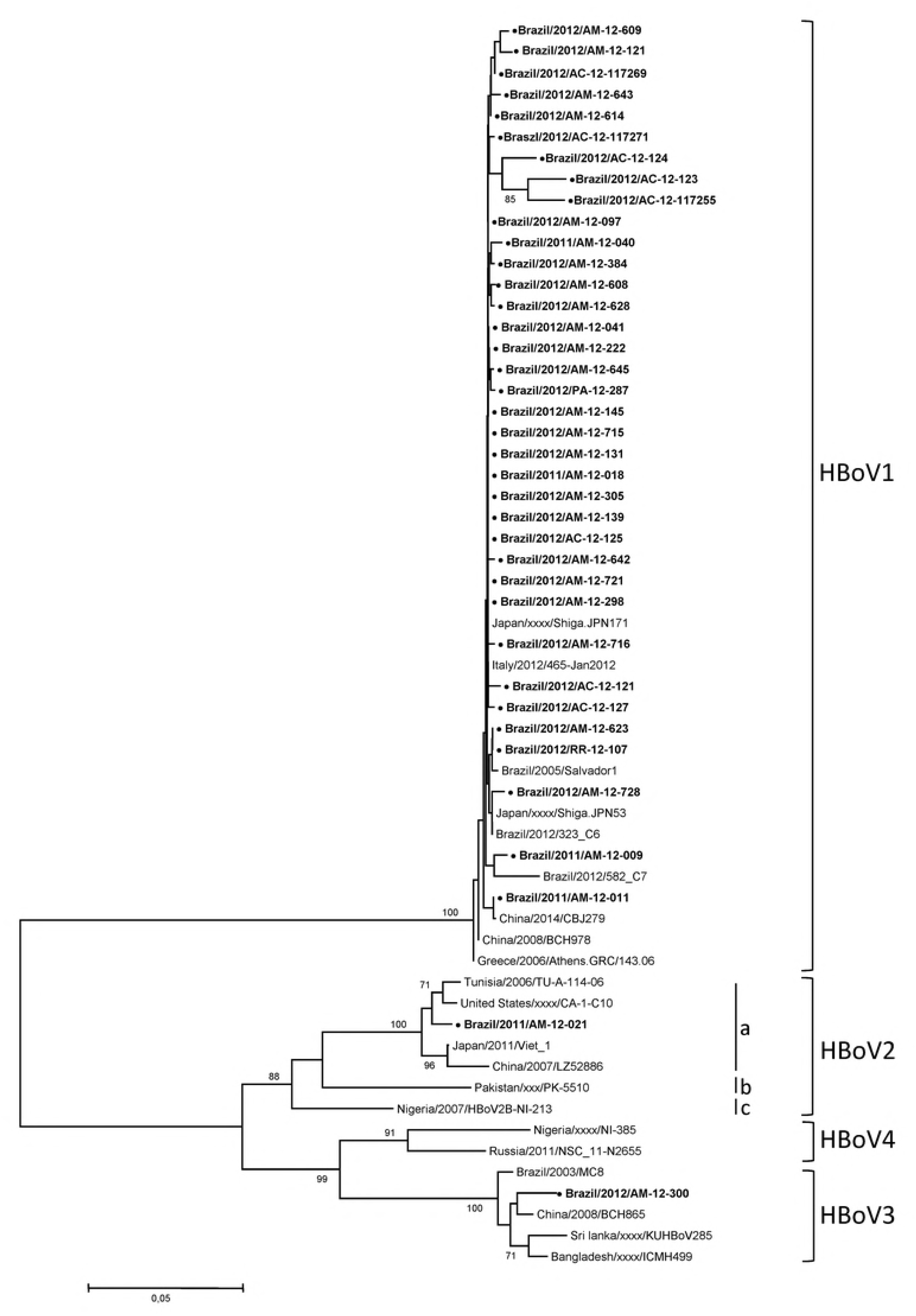
Phylogenetic analysis of VP1 protein of HBoV Brazilian strains. The level of bootstrap support is indicated at each node (values <70% were omitted) based on neighbor-joining analysis of 2000 replications. HBoV strains analyzed in this study are in bold and with circle (•).

## DISCUSSION

This study reports HBoV detection among pediatric patients with acute gastroenteritis from Brazil, is demonstrated the prevalence of HBoV and describe its dominant species. HBoV was detected in 24.0% of patients. Presents findings were higher than most of the previously reported in Brazil, including a survey with HIV-seropositive children hospitalized with acute diarrhoea, in which HBoV overall positivity rate was 14% (4,29–31).

HBoV and RVA co-infection is usually related. In the current study, this condition was observed in 50% of samples tested, a number comparable to the obtained for HBoV and diverse gastroenteritis viruses in an investigation performed in West China, between 2012 and 2013, in which the overall rate of co-detection was equivalent to 43,3% (23). In Pakistani children, 98% of HBoV-positive samples also tested positive to RVA, a higher number than observed in the present results. It is important to notice that RVA vaccine is not included in vaccination program in Pakistan, and this may be an important contributing factor to the high prevalence of RVA among infants from that population (24). On the other hand, in Brazil, where the immunization program includes RVA vaccine, HBoV and RVA association was also found, but the major co-infection rate was noted between HBoV and noroviruses (12).

As the co-infection data were limited to RVA in the present investigation, since it was the only viral agent tested other than HBoV, it was not possible to determine the relation of HBoV with other acute gastroenteritis causative virus. Another limitation consists in the absence of enough clinical data regarding to HBoV mono- and co-infections. Hence, it was not possible to stablish if there were significant differences in disease severity between these groups.

Despite HBoV-2, -3 and -4 are most associated with AGE symptoms while HBoV-1 is most related to respiratory tract diseases (Padhye et al., 2016), we identified a higher presence of HBoV-1 among HBoV-positive stool samples compared to the remaining species. Similar results were observed in Chile, where HBoV-1 was found in 14,1% of the HBoV-positive fecal specimens, between 1985 to 2010, and in other countries such as India, Thailand, Pakistan, South Korea and Australia (14,24,32–35).

An investigation performed at the same period (year of 2012) in Bahia state, Northeast Brazil, revealed, however, that 70% of HBoV-positive sequenced samples were correspondent to specie 2 (subtype 2A), while only 30% were related to HBoV-1 (12). In Southeast Brazil, a preview study has shown that 20.8% of a diarrheic group was characterized as HBoV-2, whereas 1.2% corresponded to HBoV-1 (29).

There is a current lack of data regarding to HBoV infection in Northern region of Brazil. Thus, it is difficult to compare the obtained results with other investigations about HBoV in this population and, likewise, to stablish the aspects of HBoV infection in this region. Therefore, based on the different species profile observed between distinct Brazilian regions, it is necessary to understand which peculiar characteristics from each location stimulate the propagation of diverse types of HBoV in several time spaces. Some factors as climate changes, economy and immunological situation of individuals are thought to be probable determinants of the observed particularities (36).

According to (37) the prevalence of HBoV species may fluctuate over the years, however, it was not possible to determine if the higher detection rate of HBoV-1 in Brazil was due to this factor, as only one year was evaluated on the present investigation.

Further investigations are necessary to improve the knowledge on the role of HBoV in gastrointestinal infections. Once HBoV was found with and without association to RVA, new data are important to understand the possible role of this virus in acute gastroenteritis.

## ACKNOWLEDGEMENTS

The authors would like to acknowledge the staff of State Central Laboratories (LACENs) and the technical assistance given by the laboratory personnel at the Seção de Virologia of the Instituto Evandro Chagas. The authors are also thankful to the children/mothers who agreed to participate in this study as volunteers and permitted the analysis of their relevant biological material.

## AUTHOR’S CONTRIBUTION

ABFL, KCP, JFC: performed the experiments. PSL, SFSG, DAMB, RSB: carried out molecular and the phylogenetic analyses. LSS, ABFL: analysed, interpreted data and drafted the manuscript. LSS, JDPM: provided critical review of manuscript. All authors read and approved the final manuscript.

